# Theory of mechano-chemical patterning and optimal migration in cell monolayers

**DOI:** 10.1101/2020.05.15.096479

**Authors:** Daniel Boocock, Naoya Hino, Natalia Ruzickova, Tsuyoshi Hirashima, Edouard Hannezo

## Abstract

Collective cell migration offers a rich field of study for non-equilibrium physics and cellular biology, revealing phenomena such as glassy dynamics [1], pattern formation [2] and active turbulence [3]. However, how mechanical and chemical signaling are integrated at the cellular level to give rise to such collective behaviors remains unclear. We address this by focusing on the highly conserved phenomenon of spatio-temporal waves of density [2, 4–8] and ERK/MAPK activation [9–11], which appear both *in vitro* and *in vivo* during collective cell migration and wound healing. First, we propose a biophysical theory, backed by mechanical and optogenetic perturbation experiments, showing that patterns can be quantitatively explained by a mechano-chemical coupling between three-dimensional active cellular tensions and the mechano-sensitive ERK/MAPK pathway. Next, we demonstrate how this biophysical mechanism can robustly induce migration in a desired orientation, and we determine a theoretically optimal pattern for inducing efficient collective migration fitting well with experimentally observed dynamics. We thereby provide a bridge between the biophysical origin of spatio-temporal instabilities and the design principles of robust and efficient long-ranged migration.

---

The collective dynamics of cell migration has been a topic of intense interest for biologists and physicist alike, due on the one hand to its crucial role in embryonic development, wound healing, homeostasis and cancer invasion [12, 13] and on the other hand to it being a prime experimental example of complex dynamics in out-of-equilibrium physics [14, 15]. Epithelial cells can indeed introduce local energy to the system to drive their collective migration, exerting forces on substrates which are also transmitted to neighbouring cells via adhesive complexes. Thus, epithelial monolayers are an ideal system for understanding how cell assemblies coordinate their forces and movements. Elegant *in vitro* experiments have revealed a wealth of intricate collective dynamics, ranging from long-ranged directed migration and fingering instabilities towards free edges [16, 17] to glassy dynamics [1], active turbulence [3] and spatio-temporal density waves [2, 4–8]. Conversely, biophysical studies have aimed to model these dynamics, in particular by concentrating on the mechanical and hydrodynamical properties of epithelial monolayers [2, 5–8, 18–22],

On the other hand, whether the internal state, i.e. biochemical signalling, of individual cells influences the resulting dynamics remains less explored, both theoretically and experimentally. This is in part due to the practical challenges involved in assessing protein activity live and with high temporal resolution, although recently the development of FRET biosensors [23, 24] has begun to alleviate some of these difficulties. Imaging monolayer movements together with ERK/MAPK activity has for instance revealed that the previously observed spatio-temporal waves in cell density are accompanied by corresponding waves of ERK activity (with ERK activity locally anti-correlated to cell density) [10]. Strikingly, such ERK waves were also observed in vivo during wound healing in mouse skin [25], and shown to be crucial for robust directional migration in response to a wound in *vitro* [11]. Given that the ERK/MAPK pathway is a central signalling platform, able to integrate and modulate mechanical forces [10, 11, 26–28], these experiments raise the intriguing possibility that mutual mechano-chemical feedback loops control the behaviour [5, 29, 30], although this remains to be systematically and theoretically explored in monolayers.

Here, we propose a biophysical model of collective motion in epithelial tissues, coupling the mechanics of monolayers to the mechano-sensitive dynamics of ERK/MAPK activity, both from first principles and experimental observations [11]. We uncover a mechano-chemical instability which produces complex spatio-temporal patterns of cell density, cell velocity and ERK activity at a finite temporal period and length scale. We test the model and fit all of its parameters through analysis of independent optogenetic and mechanical stretching experiments of MDCK monolayers [11], to provide parameter-free quantitative predictions of multiple features observed in experiments including the period and wavelength of instability. Next, by coupling this instability to active cell traction, we demonstrate theoretically the existence of an optimal pattern for guiding collective migration, via robust long-ranged alignment of traction forces towards free edges. Strikingly, this optimal pattern is close to the ones observed both *in vitro* in MDCK layers [10, 11] and *in vivo* in mouse skin wound healing [25]. Finally, we challenge the model through genetic and pharmacological perturbation experiments and discuss possible implications of this theory for other biological settings.

## RESULTS

Firstly, in agreement with previous studies of expanding MDCK monolayers [9–11], we confirmed that directional waves of density and ERK activity propagate away from leading edges towards the center of colonies (Fig. S1a,b). Moreover, we performed the same experiments at full twodimensional confluency (Fig. 1a-d), and found similar waves of ERK and cell density (with ERK positively correlated, and slightly lagging behind, 2D cell area), although in this case they displayed random orientation of propagation. This suggests that monolayer expansion (mediated by leading-edges) is not required for wave formation (Fig. 1a-d and Supplementary Movie 1), motivating us to first study theoretically how isotropic waves of ERK and density arise in the bulk.

**Figure 1:**
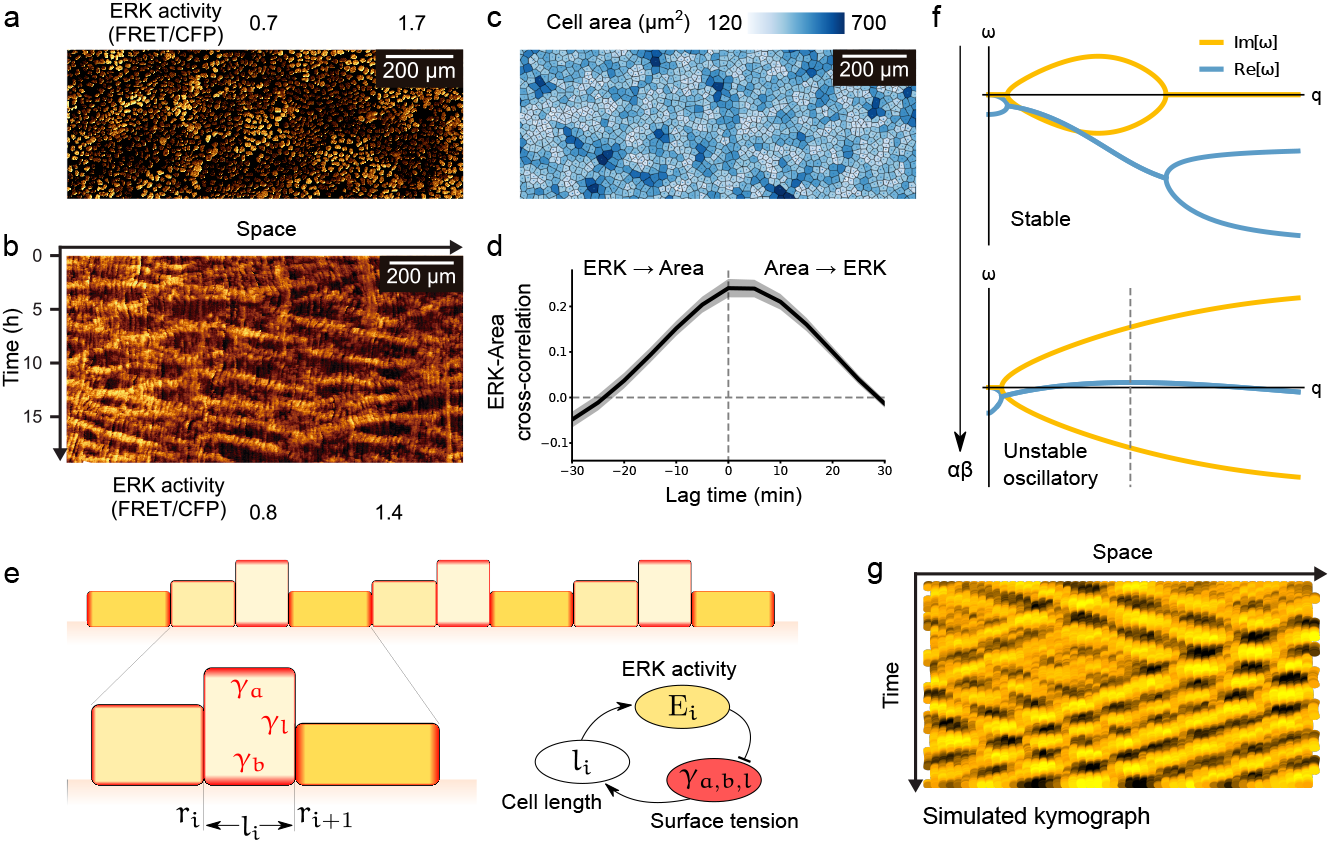
Observation and theory of spatio-temporal patterns in confluent cell monolayers. **a-c)** Confluent MDCK monolayers display ERK activity (a) and density waves (b, color-code indicates cell area), although in contrast to migrating monolayers these waves are un-directed (kymograph in panel c, see also Supplementary Movie 1). d) Cross-correlation function of cell area and ERK activity indicates a robust positive correlation between the two [10], with ERK trailing slightly area by around 3 – 5 min [11] (average of N=3 experiments - shaded areas indicate standard deviation) e) Schematic of our mechano-chemical model. ERK activation (yellow) causes actomyosin (red) remodelling, differentially affecting apical-basal and lateral tensions *γ_a,b,l_*. The dependence of ERK activation on cell length *l* completes a mechano-chemical feedback loop between ERK activation, cortical tensions and cell aspect ratio. f) Linear stability of this model (Eq. 4) reveals a finite wavelength oscillation (Re[*ω*] > 0 -dashed line) when the strength of the mechano-chemical feedback loop *αβ* exceeds a certain threshold. g) This instability is confirmed in numerical simulations of the model (kymograph).

For this, we first wrote down a minimal model of epithelial monolayer mechanics which we highlight here (see SI Text Section I for details). The three-dimensional shape of MDCK monolayers, like other epithelial tissues, is proposed to arise from a balance of forces between the active stresses generated by actomyosin cortices at the apical, lateral and basal surfaces [31–33] (resp. denoted as *γ_a_, γ_l_* and *γ_b_*). For a single cell on a flat surface, equilibrium of these forces requires the in-plane radius of a cell *l* to be equal to a ratio of active tensions, such that 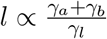 However, for confluent, heterogeneous, epithelial tissues, cell-cell junctions impose mechanical couplings between cells [35], so that the mechanical state in each cell influences the shape, density and velocity of its neighbours [36]. To describe this we consider, in a simple 1D setting, a linear chain of coupled cells with vertex positions *r_i_* (Fig. 1e) and surface tensions *γ_i_*. If a given cell contracts, it imposes a stress on its neighbours, which is resisted by cellular tensions, as well as by frictional forces with the substrate *f_i_* = −*ζv_i_* (where *ζ* is a cell-substrate friction coefficient and *v_i_* = *∂_t_r_i_* the velocity of the cell vertex *r_i_*).

At the continuum limit and at linear order, force balance in such a monolayer reads

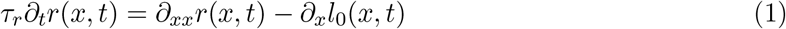

with all units normalized by the average cell length < *l* >. The first two terms are classical for overdamped chains of oscillators, and represent resp. frictional forces (with *τ_r_* = *ζ/k* the ratio of friction to cell stiffness, giving a time scale for the relaxation of mechanical stresses) and restoring forces from active cell tensions. The third term arises from the fact that each cell at position *x* can in principle have its own preferred radius *l*_0_, so that gradients of preferred radius result in cellular flows. We neglect here long-term viscoelasticity arising for instance from rearrangements, and cell division/death [34], as it occurs on time scales longer than the observed density oscillations [6].

We now seek to couple this mechanical model to the internal chemical state of the cells. Given the reported impact of MAPK/ERK on actomyosin [5, 10, 11], we assume generic couplings between ERK activity and all tensions *γ_a,b,l_*(*ERK*). Importantly, as each of these dependencies can be different, one can generically predict that ERK activity will modify the preferred radius (*l*_0_ = *l*_0_(*ERK*)) (and thus the preferred cellular density), with a delay which we denote as *τ*_l_. This is supported by recent evidence in MDCK monolayers, which showed that varying ERK activity causes relative changes in F-Actin intensity and structure in the lateral and basal surfaces [11]. Assuming linear, first-order kinetics, this effect can be described as

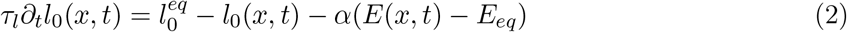

where 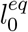 is the normal, “homeostatic” rest length (which sets the length scale of the problem) corresponding to steady-state ERK activity *E_eq_*, and *α* is the coefficient of coupling between ERK activity and preferred shape changes. In principle *α* can have any sign, however optogenetic activation of ERK in MDCK monolayers causes cellular contraction [11], suggesting that *α* > 0 in this system.

Finally, given the reported mechano-sensitivity of ERK and its dependency on cell length (*∂_x_r*, inversely proportional to cell density) [11], a minimal equation for ERK activity, again assuming generic first-order couplings, reads

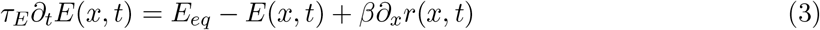

with the time scale *τ_E_* indicating how quickly ERK reacts to mechanical deformations, and *β* determining the strength of coupling between cell length and ERK levels. Live-measurements of ERK activity demonstrate that stretching MDCK monolayers results in an activation of *ERK* [11], suggesting that *β* > 0 in this system (see SI Text Section ID 1 for further details).

We now re-define variables *E* = *E* — *E_eq_* and 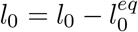 so that we only look at perturbations around steady state, and the full model reads

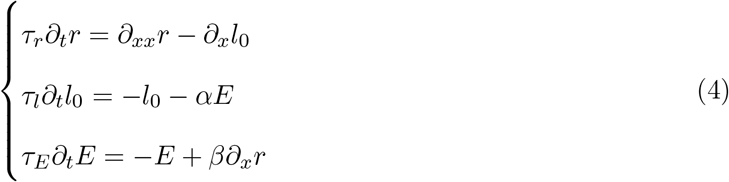

We perform a linear stability analysis to test whether this minimal mechano-chemical model reproduces the features of density/ERK patterns observed in epithelial monolayers. This yields the following dispersion relation for patterns of temporal frequency *w* and spatial frequency *q*:

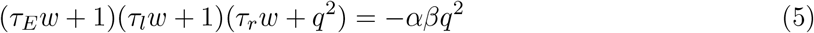

We see that there are only three independent quantities in this problem: two ratios of timescales (*τ_E_/τ* and *τ_r_/τ_l_*), as well as the product of the two coupling coefficients *αβ*, which quantifies the global strength of the ERK-density mechano-chemical feedback loop. Examining the evolution of the dispersion relation *w*(*q*) reveals a critical threshold for *αβ* above which an instability occurs (see SI Text Section IB), characterised by a finite length scale and non-zero imaginary frequency indicative of stable temporal oscillations (Fig. 1f). This means that complex spatio-temporal patterns of ERK and density are expected to arise with lengthscale

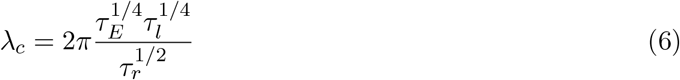

and temporal frequency

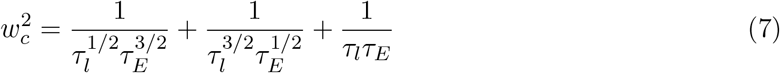

which is confirmed in numerical simulations (Fig. 1g and Fig. S2a,b). Since the frequency of the emergent oscillation depends only on the timescales *τ_l_* and *τ_E_*, we can infer that the instability arises from local activator-inhibitor dynamics between the preferred cellular radius and ERK activity. The remaining timescale *τ_r_*, for the magnitude of substrate friction/mechanical relaxation, determines how far local deformations propagate over one period of oscillation, which explains the finite length scale of the instability. In contrast to previous approaches relying on polar migration forces [5–8], this minimal model requires only scalar active terms to produce patterns of density, velocity and ERK at finite length and time scales. We also note that density waves are abolished by inhibition [5], but also overactivation of ERK (Fig. S1), which is in full agreement with the predictions of our model (Fig. S2c), and shows that ERK oscillations are not merely a byproduct of an upstream density oscillation (as observed for YAP oscillations [7]), but rather a core element of the dynamics.

Importantly, Eqs. 6 and 7 make predictions that depend only on macroscopic parameters, i.e. the three time scales *τ_r_, τ_E_* and *τ_l_*, which we now seek to infer from mechanical stretching and optogenetic activation experiments in MDCK monolayers [11]. Firstly, mechanical stretching accompanied by live-reporting of ERK activity (Fig. 2a) offers the ideal platform to measure *τ_E_* since it is the timescale of ERK activation post-stretch. Comparing Eq. 3 to the data of Ref. [11] reveals good fits (Fig. 2b and Fig. S4a), which validates our hypothesis of first-order kinetics and dependency of ERK upon cell density, and allows us to infer *τ_E_* = 4 — 8 min (we also took into account longer-term desensitization of ERK, although this has little impact on the results, see SI Text Section ID1 and Fig. S3c,d for details). Secondly, optogenetic activation of ERK and observation of subsequent cellular contraction [11] (Fig. 2c) provides a dataset from which to infer *τ_r_* and *τ_l_*, as well as an opportunity to validate the mechanical model (Eqs. 1 and 2) for externally imposed ERK dynamics. When a step function in ERK activation is applied, the model predicts cellular shrinking at timescale *τ_l_* beginning at the boundary between high and low ERK, where forces are unbalanced, and diffusing into the monolayer at timescale *τ_r_* (see SI Text Section IVB). We thus tracked cells to follow the spatio-temporal dynamics of monolayer contraction *r(x, t)* (raw data from [11]). Fitting this dataset to our analytical prediction (Eq. S39) reveals excellent qualitative and quantitative agreement, which allows us to infer *τ_l_* = 100 — 140 min and *τ_r_* = 4 — 11 min (Fig. 2d-f and Fig. S4b-d). Strikingly, these estimates predict a typical length scale *λ_c_* and temporal period *T_c_* of density/ERK patterns at the onset of instability

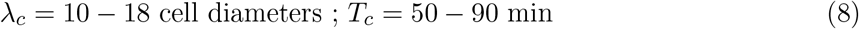

which describes well, and in a parameter-free manner, the dynamics observed experimentally both in confluent and migrating MDCK layers (Fig. 1a-d, Fig. S1a,b and Supplementary Movie 1). Furthermore, the model predicts ERK waves to trail behind density waves by a characteristic time-scale of *τ_E_*, in agreement with our data (3 — 5 min from Fig. 1d).

**Figure 2:**
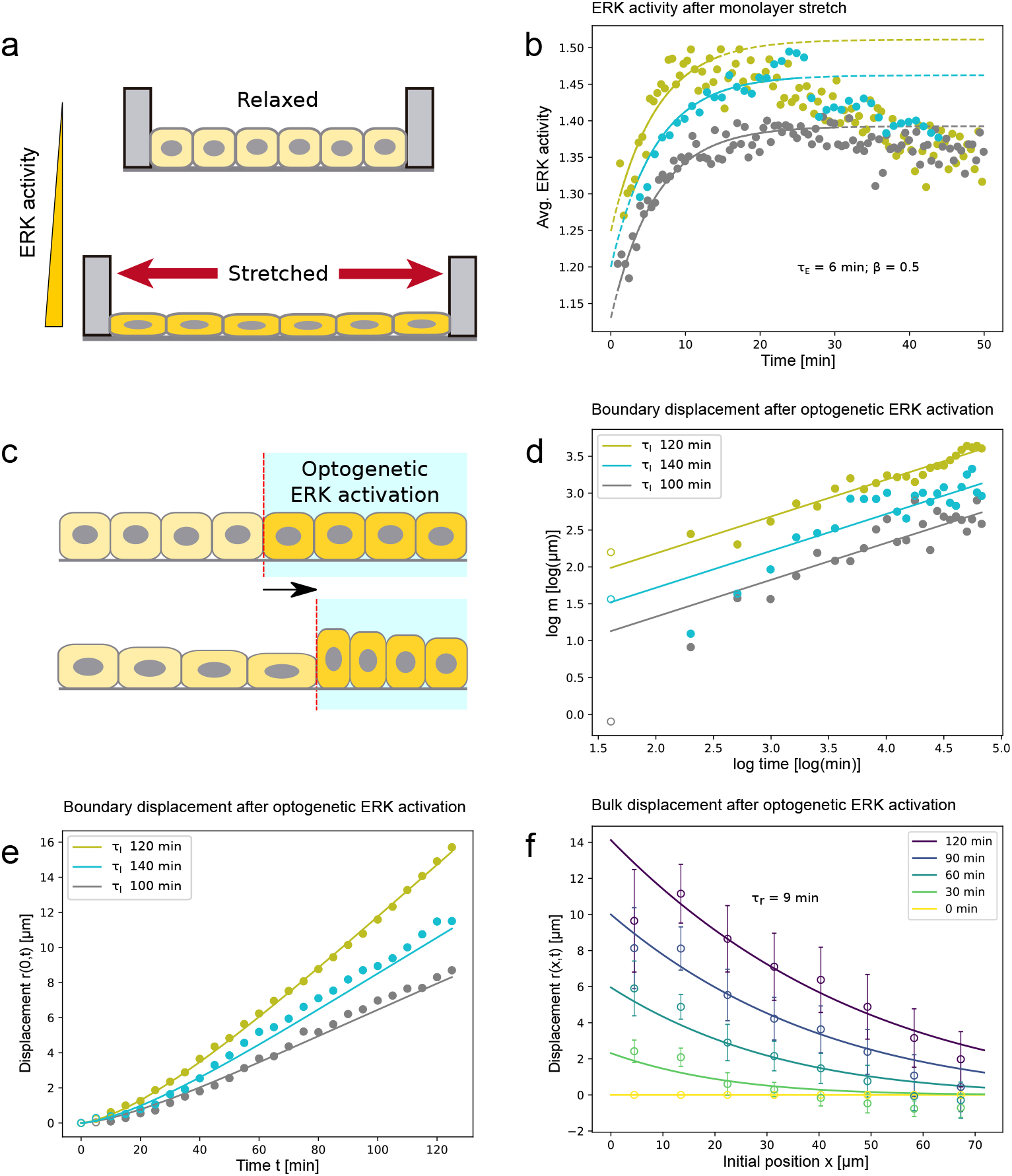
Parameter-fitting using mechanical and optogenetic perturbation experiments. a) Response of ERK activity to a 50% uni-axial stretch (N=3, from Ref. [11]). b) Eq. 3 provides an excellent fit for the data (dots, each color shows an independent experiment), from which we extract *τ_E_* = 4 — 8min (and *β* = 0.5 — 0.6). Solid lines indicate the model fits. c) Optogenetic ERK activation in patch of cells cause cellular contraction from the boundary (N=3, from Ref. [11]). d-f) Our model provides a good fit for the displacement of the boundary during contraction (panels d,e: each color shows an independent experiment) as well as the full spatio-temporal evolution of the cell displacement field *r*(*x, t*) (panel f) upon ERK activation at *t* = 0 (see SI Text Section IVB), allowing us to extract *τ_l_* = 100 — 140min and *τ_r_* =4 — 11min. Error bars show average and standard error from the three repeats.

Although our proposed mechanism is sufficient to explain isotropic patterns in confluent tissues, we still need to explain how symmetry is broken to allow for unidirectional waves and long-range cell polarity in migrating monolayers with a free boundary (Fig. 3a) [11]. For this, we couple our model to cell polarity 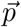 (sketched in Fig. 3b), generated in MDCK layers by polarized lamellipodia [6, 40], which enters as an additional traction force in Eq. 1 (see Eq. S14). It has been reported, in particular, that cells polarize in response to tension transmitted through cell-cell junctions [6, 15, 39], so that the simplest equation for traction force orientation reads

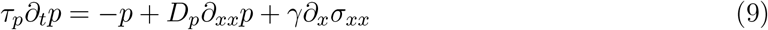

where *τ_p_* is a time scale of relaxation, *D_p_* is a diffusion coefficient representing possible polarity alignment between neighboring cells [15] and *γ* > 0 is a coupling strength between polarity and gradients of cellular stresses, *∂_x_σ_xx_* = — *∂_x_*(*∂_x_r* — *l*_0_) (see SI Text Section ID3 for a discussion of alternative couplings, e.g. with velocity [18]).

**Figure 3:**
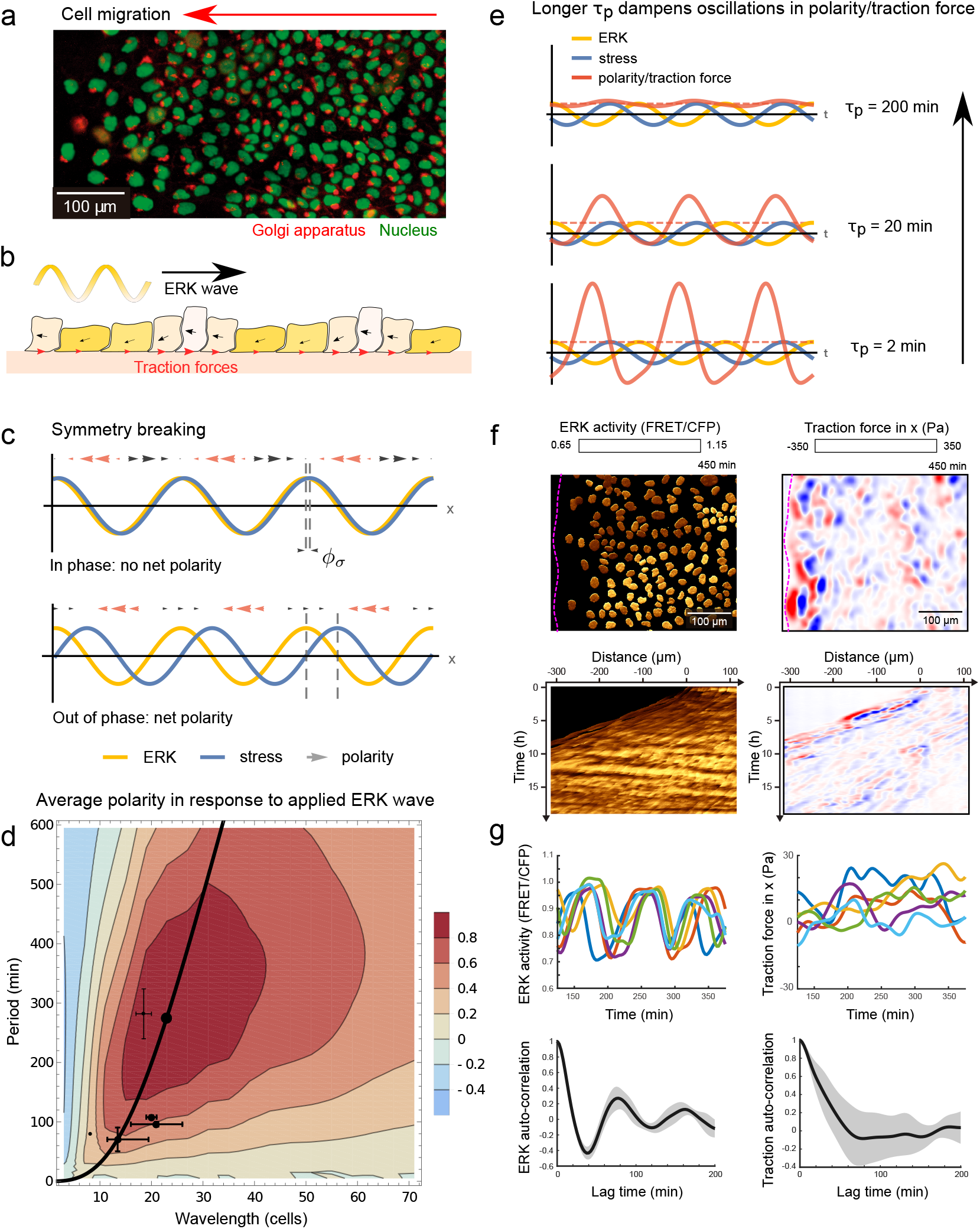
Response of cell polarity to an applied ERK wave and comparison with traction force data. a) Migrating MDCK monolayers polarize towards a free edge as visualized by the orientation the Golgi apparatus (red) relative to the nucleus (green) (see Ref. [11]). b) Schematic of our mechano-chemical model extended to include active cell migration/polarity. c) If polarity follows gradients of stresses, monolayers can exploit phase differences between ERK and stress to produce net polarization from symmetric waves. d) Quantitative phase diagram of the average cell movement in response to applied ERK waves, showing an optimal period and wavelength for effective migration (positive values in red indicate polarization counter to ERK waves). Data points are previously reported values for wavelength and period in MDCK layers [8, 10, 11] and mouse skin [25] (see SI Text Section IC for details). Importantly, the waves permissible by our biophysical model (black curve for varying *τ_E_*) produce only positive net polarization, thus providing robustness to the migration response. e) Oscillations in polarity/traction force are dampened for longer timescales of polarization *τ_p_* whereas average polarization (dashed line) is unaffected. f) ERK activity and corresponding traction forces measured in an expanding monolayer. Dashed lines indicate the leading edge of the monolayer. Kymographs show that the multi-cellular waves present in ERK are absent in traction forces. g) Single cell traces (5 representative cells, color-coded) of ERK activity show strong oscillatory component (auto-correlation function), whereas corresponding local traction forces show a much weaker oscillatory component (as predicted in panel e). Shaded areas: standard deviation.

Before studying the fully coupled system, we first explore the limiting case of externally driven ERK dynamics (Fig. 3b). Although the couplings in our model allow ERK waves to translate into changes in polarity, a solely linear theory cannot break symmetry to produce unidirectional polarization. This is similar to the “back-of-the-wave” paradox encountered in the collective migration of *Dictyostelium* [37, 38]. However, we reasoned that non-linearities (combined with phase differences between ERK and mechanics) can allow for symmetry breaking, for instance if the stress-polarity coupling is modulated by ERK activity (γ = γ(E)). This is consistent with recent experiments where ERK activation (resp. inhibition) was shown to decrease (resp. increase) mean traction forces in MDCK layers [11], implying moreover that *γ* is a decreasing function of *E*. This could provide a generic mechanism to take advantage of phase differences between ERK activity and me-chanical stress, to allow cells to respond strongly to the positive gradients of approaching mechanical waves, while being largely insensitive to the next negative gradient (see Fig. 3c for a sketch). When we analyse this we find that a maximum global polarization occurs when stress lags ERK by 3*π*/4, which occurs only at a single optimum of the wavelength λ and temporal period *T* of the applied ERK wave (Fig. 3d and SI Text Section IC).

Using the timescales inferred above allows us to predict the values of this optimum (Fig. 3d), which we find to be λ ≈ 15 — 40 cells and *T* ≈ 2 — 8 hours (stochastic simulations of the full model with ERK desensitization, or slightly longer for the analytical criterion, see SI Text Section I D 1 and Fig. S5). Strikingly, this is close to the natural waves observed both in MDCK layers (λ ≈ 20 cells, *T* ≈ 1 h in our data and Ref. [10, 11], λ ≈ 20 cells, *T* ≈ 4.7 h in [8]) and mouse skin (λ ≈ 8 cells and *T* ≈ 1.3 h [25]), suggesting that these systems might be tuned for optimal migration. This is also in agreement with optogenetic experiments which applied waves of a given wavelength, but varying speed (i.e. period), to MDCK monolayers, demonstrating the existence of an optimal speed for cell movement [10] (≈ 2*μ*m/min in both data and our model). Furthermore, although any wave characteristic is possible in optogenetic experiments, the mechanism for spontaneous pattern formation which we described above permits only waves which translate into net polarization in the direction opposite to the propagation of ERK (Fig. 3d). In other words, the system robustly directs migration counter to ERK waves for any model parameters.

We also investigated how polarity is predicted to evolve spatio-temporally in response to an applied ERK wave (see Eq. S20). Although the average polarity shown in Fig. 3d is independent of the polarization relaxation time scale *τ_p_*, the amplitude of polarity oscillations around the average depends critically on it, with longer values dampening the amplitude of the oscillations (Fig. 3e). Interestingly, experimentally measured values (*τ_p_* ≈ 15 — 40 min [6, 11, 39, 40]) were sufficient to abolish negative contributions to polarity in response to an applied wave. This provides an explanation for the seemingly paradoxical observation that migration polarity displays near-constant long-ranged order [11] despite the presence of spatio-temporal oscillations in stress [2]. To further test whether this is also the case for the long-range orientation of traction forces, we measured traction forces profiles in a migrating monolayer together with live-ERK activity, and confirmed that oscillations in traction forces are weak compared to the strongly periodic ERK oscillations (Fig. 3f,g and Supplementary Movie 2).

Next, we explored the full interplay between mechano-chemical ERK patterns and active migration forces, which is the typical scenario of monolayer expansion in the presence of free boundaries. Simulations in 1D consisted of Eq. S14, supplemented with boundary conditions at the free edge (specifically *p = p_b_*) to model the presence of leader cells [16, 17] (see SI Text Section II). We found that simulations in the presence of bulk polarity displayed ERK waves robustly propagating backwards from the free edge (Fig. 4a), which gradually polarized cells in bulk, leading to long-range polar order (Fig. S6a) (as in the response to a driven ERK wave). This recapitulated well the experimentally observed dynamics (Fig. 4a and Ref. [11]). Conversely, for strong leader cells and comparatively low bulk polarity, the model predicts waves in the reverse direction (center to edge) (Fig. S6c), suggesting a key role for stress-polarity coupling in determining wave directionality.

**Figure 4:**
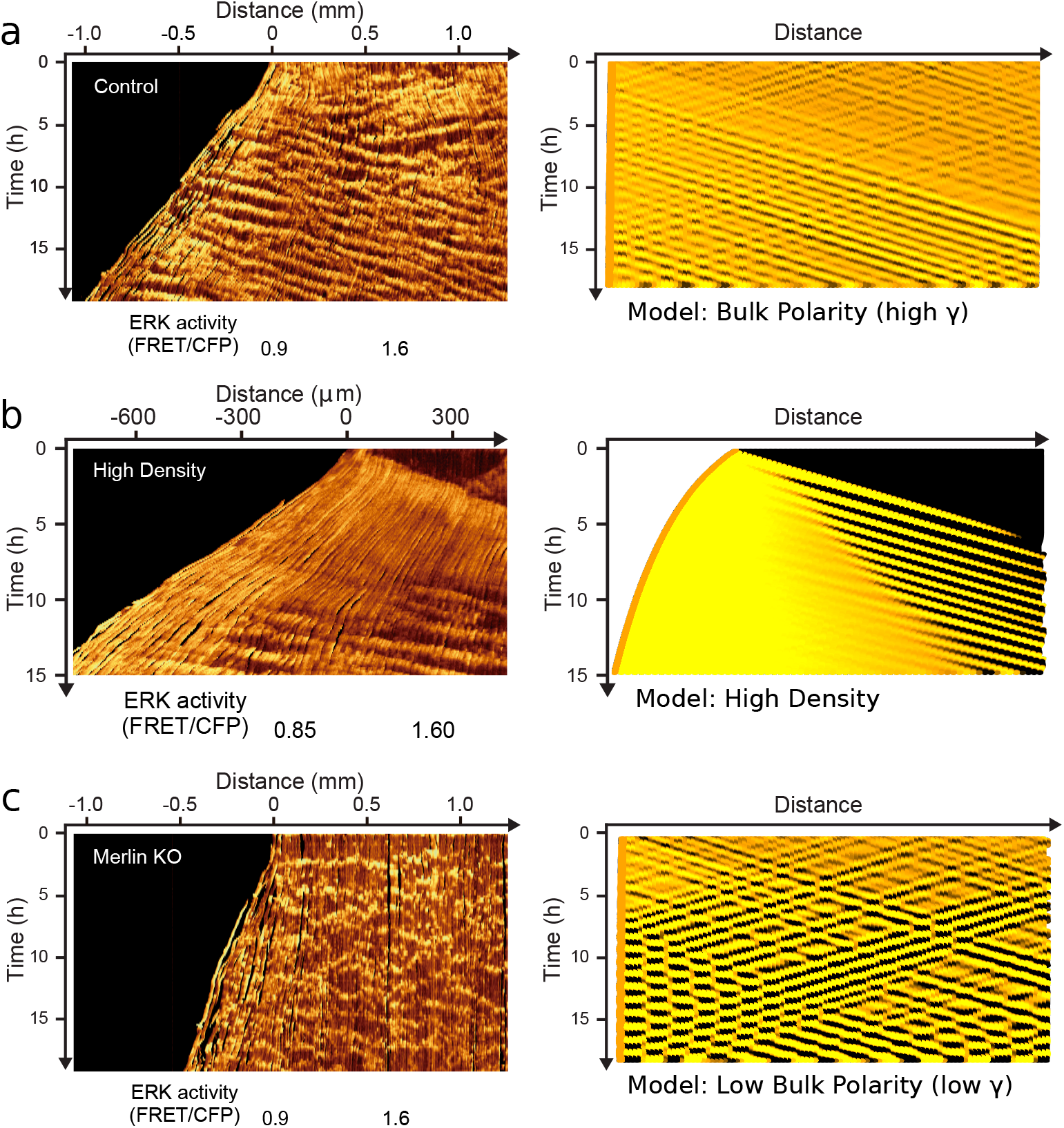
The model predicts the spatio-temporal dynamics of expanding monolayers in a variety of experimental settings. a) Using the parameters fitted above, together with a boundary condition of leader cells (SI Text Section II), the model (right) reproduces the transition from initially random to unidirectional ERK wave propagation seen in data for expanding monolayers (left). b) The model also reproduces the effect of increasing cell density, with initially low ERK activity everywhere in the monolayer, followed by a tidal wave [10] of ERK activity (which remains high in the front as waves start appearing in the back). c) Merlin knockouts (left) designed to inhibit polarization and active migration. ERK waves are still present but become randomly orientated, matching the model predicted for weaker stress-polarity coupling (*γ* ≈ 0, right).

Although multiple experimental set-ups produce density waves propagating backward from the edge [6, 11], some groups report more complex dynamics such as a front of uniformly high ERK propagating from the boundary [9, 10]. Interestingly, our model predicts this behavior for cell monolayers cultured at initially higher density (uniform cell length smaller than the preferred length, i.e. *l* < *l*_0_, as initial condition), where the first stages of expansion are dominated by mechanical relaxation (as reported in Ref. [41]) which triggers a low density/high ERK front propagating away from the boundary. To test this prediction we cultured cells at higher density before release, and found good qualitative agreement (Fig. 4b and Supplementary Movie 3), arguing that cellular density could explain the different ERK dynamics observed in previous experiments [10, 11].

Finally, we sought to test the key model prediction that stress-polarity couplings are not the origin of mechano-chemical waves, but rather cause symmetry breaking (i.e. unidirectional wave propagation in response to a free-edge) of existing mechano-chemical patterns. For this, we examined ERK dynamics in Merlin knock-out cells, as Merlin has been shown to be a key link between cell-cell mediated force transmission and Rac1-driven polarized lamellipodia formation in MDCK monolayers [40]. We thus expect the coupling coefficient 7 to be drastically reduced in this condition (*γ* ≈ 0). Strikingly, Merlin KO monolayers still displayed prominent ERK waves, with indistinguishable wavelength and time period as controls (Fig. S6d and Supplementary Movie 4), but directional ERK wave propagation was lost (Fig. 4c and Fig. S6b), in agreement with the model.

## DISCUSSION

Altogether, we provide a theoretical framework for the emergence of complex mechano-chemical patterns and long-ranged coordinated migration in MDCK monolayers. We show that spatiotemporal waves of cellular density and ERK activity arise from a scalar active matter instability, which occurs generically in the presence of delayed mutual feedbacks between tissue mechanics and mechano-sensitive ERK activity [11]. Quantifying datasets from optogenetical and mechanical perturbation experiments [11], we tested the model and extracted the key time scales of the system, providing parameter-free predictions for the appearance, wavelength and period of ERK/density patterns, which fits well with previously published data, as well as our own. In contrast to previous proposal [5–8, 20–22], our results suggest that interplays between stresses and migratory forces are not the driver of the instability itself, but are instead required to orient ERK/density waves uni-directionally and establish long-range polar order. Remarkably, we show that this mechanism robustly induces polarization in a preferred direction and that biophysical parameters are tuned for a close to optimal migration response.

Our theory thus provides a bridge between the biophysical origin of spatio-temporal patterns and the design principles underlying collective migration, with ERK playing the central role of integrating chemical and mechanical signals. Integrating other biochemical networks within this framework, such as RhoGTPase, which have been proposed to contribute to instabilities [42], would be an interesting future direction and could enrich the dynamics even further. There are also other patterning phenomena for which we could test and extend our theoretical model. For instance, tissue-scale mechano-chemical waves have also been observed to drive morphogenesis in multiple developmental settings, such as tracheal [43] and endoderm [44] morphogenesis in *Drosophila.* In the former case, waves arise from a positive biochemical feedback loop from EGFR-ERK signalling between neighbouring cells, leading to a relay mechanism and wave propagation that translate to a wave of MyosinII cable formation. In the latter case, wave propagation relies on a mechano-sensitive feedback where MyosinII-generated stresses in a cell activate MyosinII in the neighbour. Cellular-level temporal oscillations in the area of *Drosophila* amnioserosa cells have also been proposed to arise via a theory integrating tensile stresses and myosin activity [36].

Beyond biophysical mechanisms relating to pattern formation, the down-stream effects (e.g. on collective cellular invasion in cancer or stem cell proliferation/differentiation) of multiple signalling pathways have been shown to depend not only on average biochemical levels, but also on their spatio-temporal dynamics [28, 45, 46]. This begs the more general question of how spatio-temporal patterns are translated into robust cellular responses in different settings. Considering together the biophysical origin and design principles underlying these behaviours could prove a useful strategy for furthering our understanding of these process.

## Acknowledgments

The authors would like to thank Gasper Tkacik and all the members of the Hannezo and Hirashima groups for useful discussions, Xavier Trepat for help on traction force microscopy, and Michiyuki Matsuda for use of facility. EH acknowledges a grant from the Austrian Science Fund (FWF) (P 31639). TH acknowledges a grant from JST, PRESTO (JPMJPR1949). This project has received funding from the European Union’s Horizon 2020 research and innovation programme under the Marie Skodowska-Curie Grant Agreement No. 665385 (to DB), from JSPS KAKENHI Grant Numbers 17J02107 to NH, and from the SPIRITS 2018 of Kyoto University to EH and TH.

## Additional information

Additional details on the analytical model, numerical simulations and experimental designs are provided in SI Text. The data and code that support the plots within this paper and other findings of this study are available from the corresponding authors upon request.

## Author Contributions

Supervision and project conceptualization: TH and EH. Theory and simulations: DB and EH. Data analysis: DB, NH, NR, TH and EH. Experiments: NH and TH. Manuscript writing: DB, TH and EH, with input from all authors.

