## Supplementary material for "Theory of mechano-chemical patterning and optimal migration in cell monolayers": SI Figures and Text

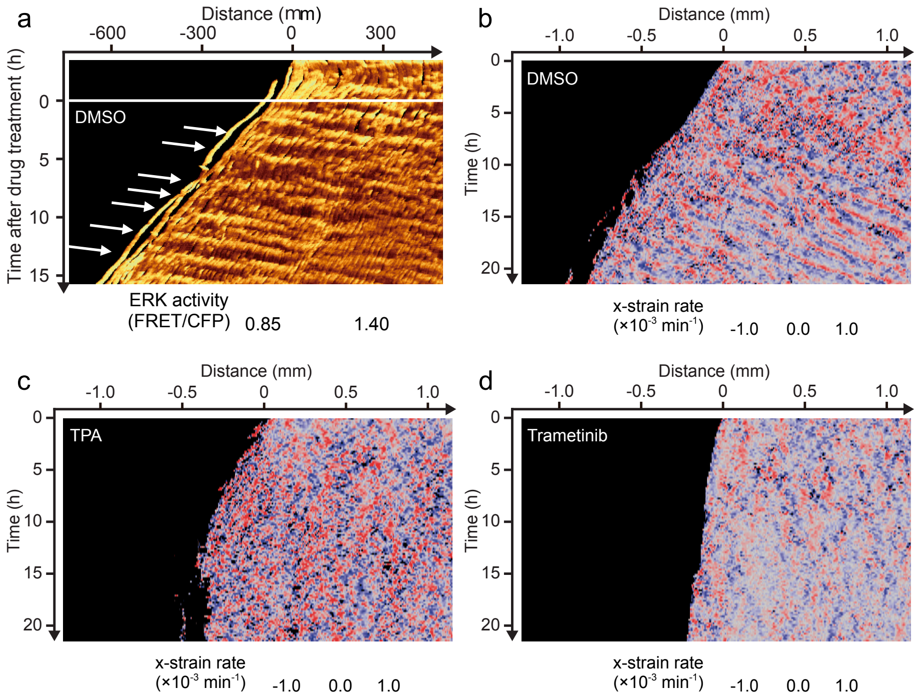


Figure S1. **Over-activation or inhibition of ERK causes loss of density waves.** a,b) DMSO controls showing WT behaviour of ERK-density waves. c,d) Drug treatments to overactivate (TPA) or inhibit (Trametinib) ERK cause loss of density waves suggesting that ERK is a core part of the mechano-chemical instability and not just a downstream component.

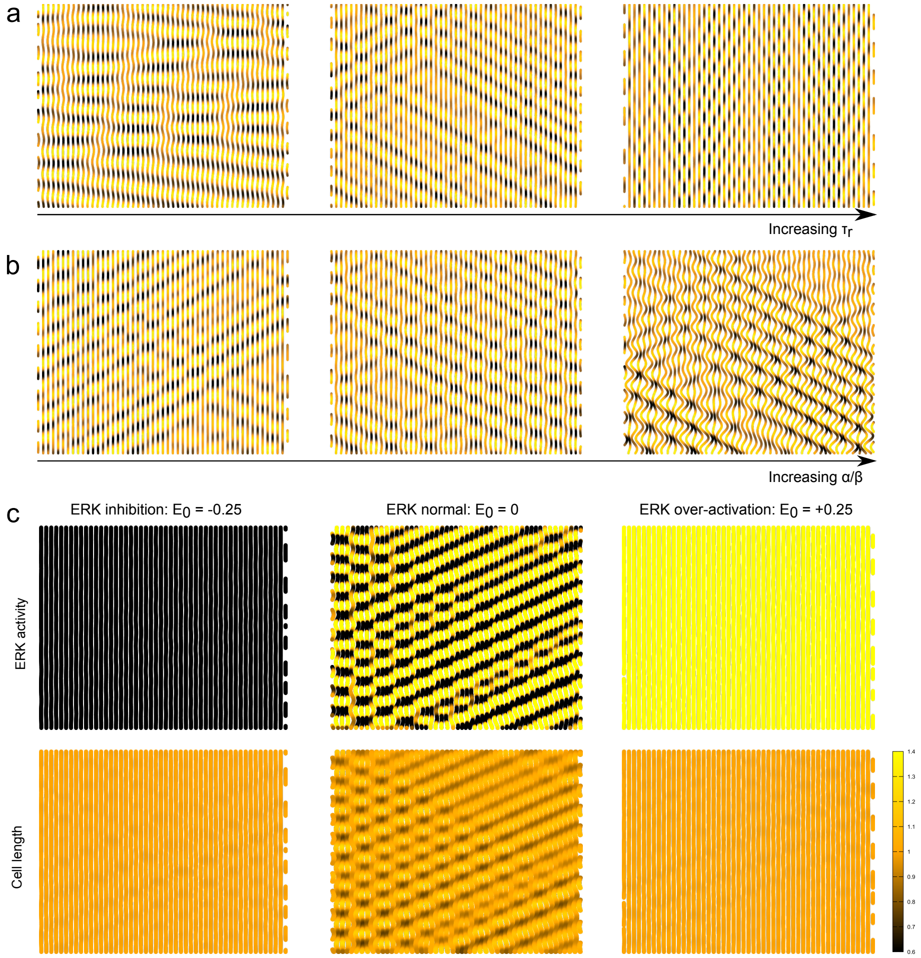


Figure S2. **Effect of varying parameters on simulated dynamics of confluent tissue.** a) Increasing the timescale of mechanical relaxation $\tau_r = \zeta/k$ (i.e. the ratio of substrate friction to cell stiffness) shortens the wavelength of the instability without changing the period. b) Increasing the ratio of mechano-chemical coupling constants $\alpha/\beta$ results in larger cell deformation without affecting the wavelength or period of the instability. All simulations use $\alpha\beta = 1.2 \times (\alpha\beta)_{\text{crit}}$. c) Waves of both ERK and density are suppressed by either inhibition or over-activation of ERK (See Eq. S31).

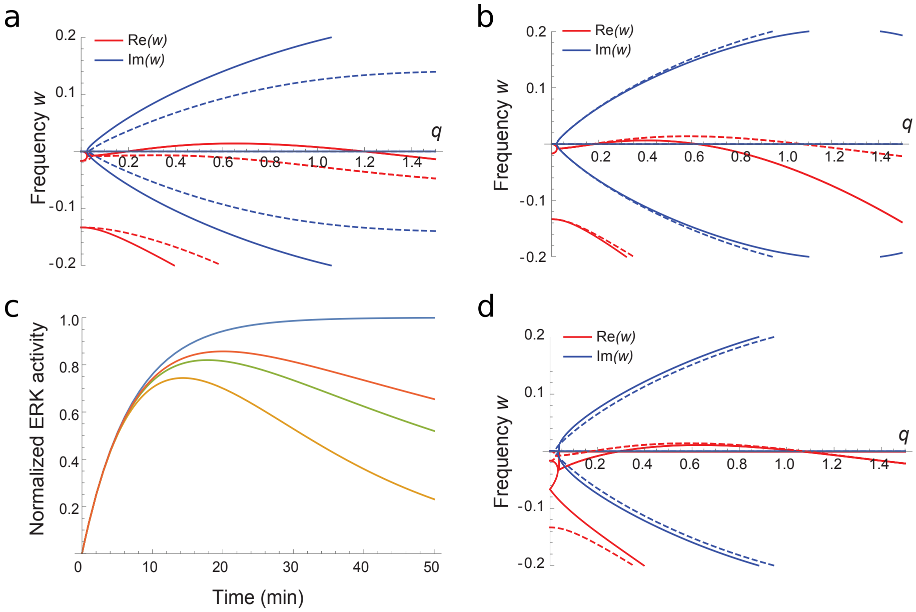


Figure S3. **Effect of different couplings on pattern formation.** a) Dispersion relation as a function of wavevector $q$ for the model presented in the main text (Eq. 4) with values of $\alpha\beta$ above (solid) and below (dashed) the threshold for instability (real part of $\omega$ in red, imaginary part in blue - positive values of the real part of $\omega$ indicate instability). b) Little changes qualitatively to this picture when we include diffusion of ERK at a realistic rate D = 1 cell$^2$/min (dashed lines). c) We can also account for the ERK desensitization (Eq. S24) seen in cell stretch (Fig. 2b) and optogenetic perturbation experiments (see [11]) (plotted $\tau_d = 30$ min (yellow), 60 min (green) 90 min (red) and $\tau_d \to \infty$ (blue)). d) But for realistic timescales this has little effect on parameter estimates or on the dispersion relation governing emergent patterns (dashed lines $\tau_d = 60$ min, solid lines $\tau_d \to \infty$).

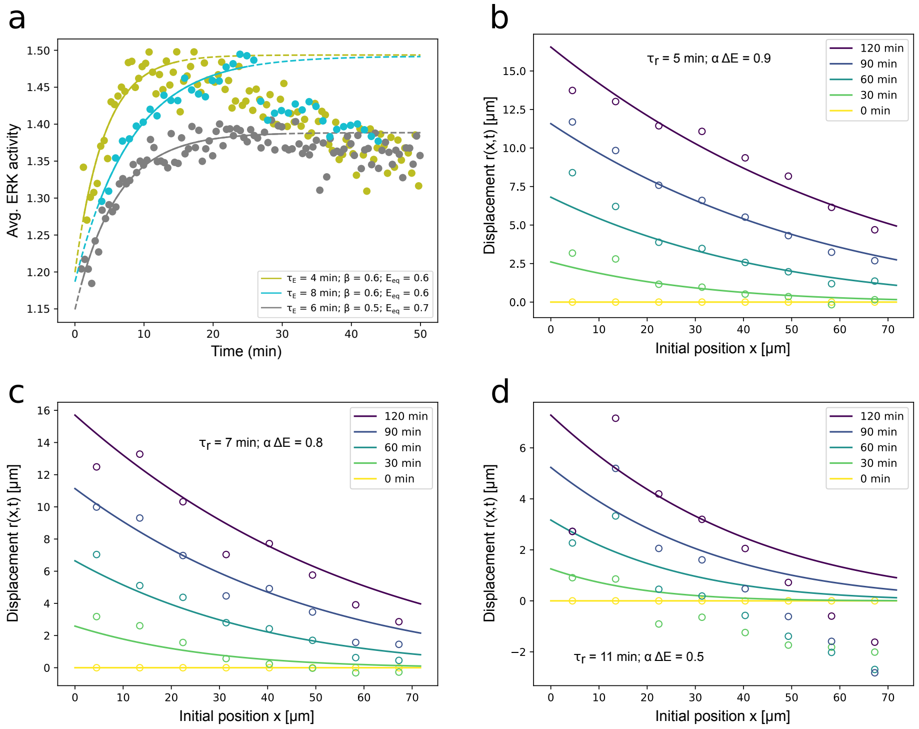


Figure S4. **Individual fits of repeats from cell stretch and optogenetic activation experiments.** a) Individual fits of the three independent repeats of cell stretch collectively fit in Fig. 2b. b-d) Individual fits of the three independent repeats for bulk displacement after ERK activation that are averaged and collectively fit in Fig. 2f. In d) displacements are distorted due to the coincidence of a natural contraction wave pre-existing the optogenetic activation of ERK and consequently data is more difficult to fit with both $\tau_r$ and $\alpha\Delta E$ as free parameters. We therefore left $\tau_r$ free but fixed $\alpha\Delta E = 0.5$ using a steady-state assumption on Eq. S34 and observation of roughly 50% difference in final cell size either side of the optogenetic boundary.

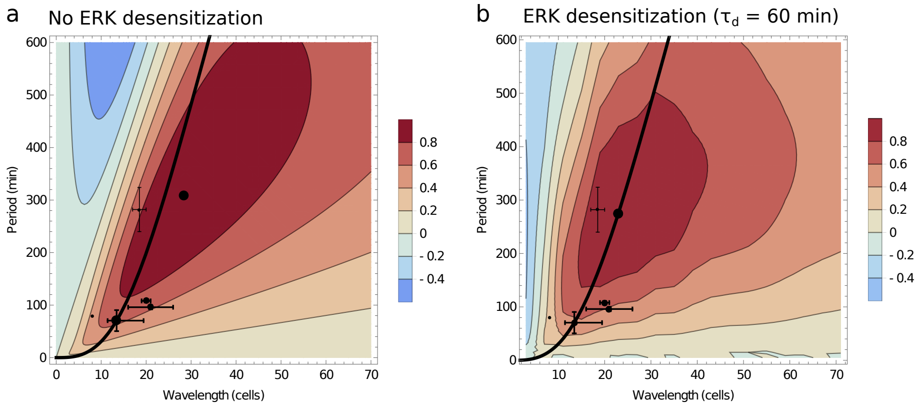


Figure S5. **Effect of ERK desensitization on optimal polarity.** a-b) ERK desensitization (Eq. S24) at realistic timescales ($\tau_d = 60$ min) decreases the wavelengths and periods of near-optimal waves. The absolute optimum moves from $\lambda = 28$ cells $T = 310$ min to $\lambda = 23$ cells $T = 275$ min. See SI Text Section I C for details of data points. (Note that b is a repeat of Fig. 2d in the main text.)

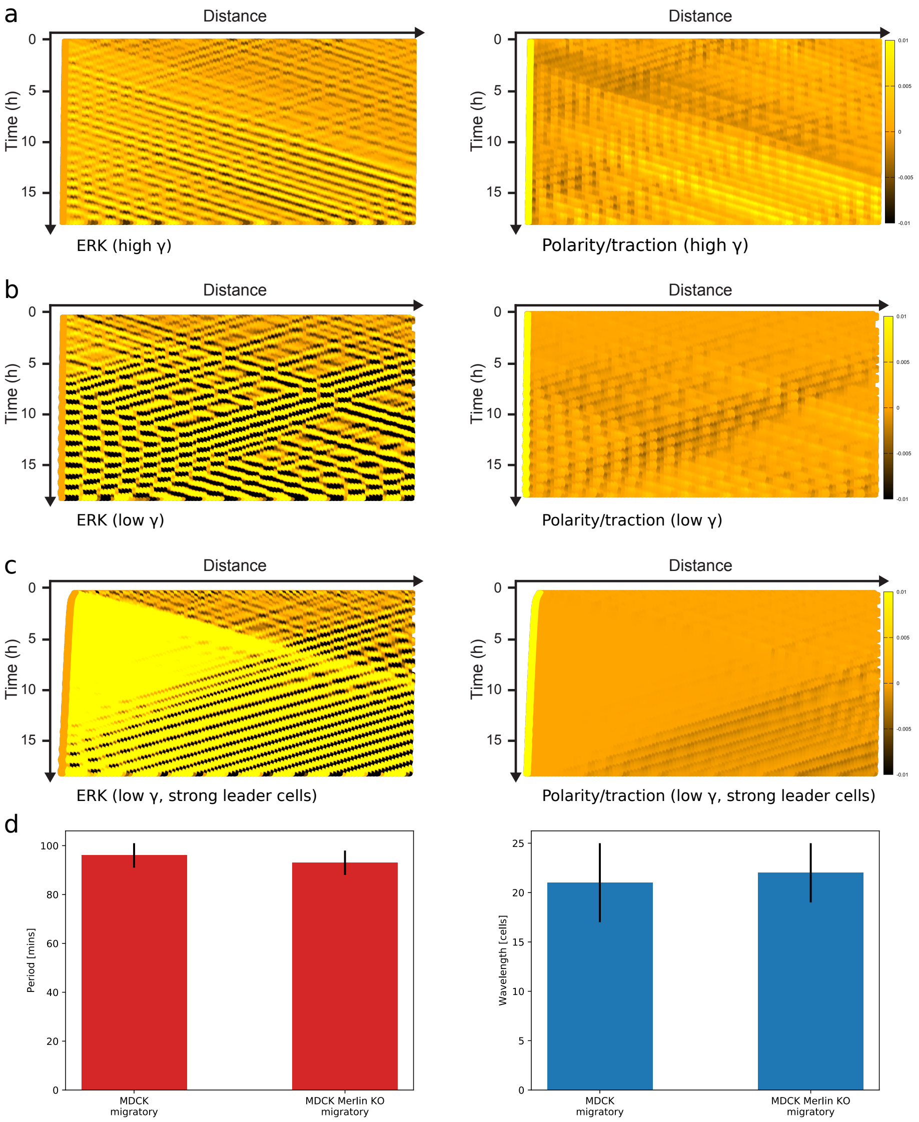


Figure S6. **Unidirectional waves and long-range polar order require stress-polarity coupling and active migration.** a) In wild-type simulations (Eq. S14), emergent ERK/density waves orient away from the edge and induce a counter polarization/active migration in the bulk. b) When stress-polarity coupling is weak waves still emerge but do not induce strong enough polarization/active migration to allow for unidirectional orientation and long-range polar order. c) Alternatively, when weak stress-polarity coupling is accompanied by strong leader cells (2.5 times stronger polarity at leading edge) waves propagate towards the migrating front and bulk polarity is established in the opposite direction. (Note that a and b correspond to Fig. 3a and c of the main text.) d) Waves with the same wavelength ($p=0.7791$, Two-sample t-test) and period ($p=0.7780$, Two-sample t-test) emerge in Merlin KOs and wild-type suggesting that active migration is not driving the instability. Error bars indicate standard deviation.

**Supplementary Information: Theory of mechano-chemical patterning and optimal migration in cell monolayers**

### I. MODEL DERIVATION

#### A. Equilibrium of cellular shape

We first start by considering the equilibrium of a single cell, under differential tensions on its apical, lateral and basal areas (resp denoted $\gamma_a$, $\gamma_l$ and $\gamma_b$). The energy of a single cell of height $h$ and characteristic length $l$ then reads [1–6]:

$$U = \gamma_l h l + (\gamma_b + \gamma_a) l^2 \tag{S1}$$

Because the 3D cellular shapes that we are modelling are restricted to a flat substrate, we need only consider stable shapes with zero average spontaneous curvature such that apical and basal areas are equal. Moreover, epithelial cells may be considered incompressible, with constant volume $V_0 = hl^2$ on the time scales considered here (minutes to hour) and given the small forces generated by actomyosin structures (compared to osmotic forces) [7]. Minimizing the energy with respect to $l$ provides the equilibrium shape of a given cell $l_0 = (\frac{V_0}{2}\frac{\gamma_l}{\gamma_a+\gamma_l})^{1/3}$, and expanding up to 2nd order around this minimum energy yields:

$$U(l) = U(l_0) + (l - l_0)U'(l_0) + \frac{(l - l_0)^2}{2!}U''(l_0) + ... \tag{S2}$$

$$= U(l_0) + \frac{(l - l_0)^2}{2!}U''(l_0) + ... \tag{S3}$$

$$\approx U(l_0) + k(l - l_0)^2, \tag{S4}$$

which is the potential of a harmonic oscillator, valid for small perturbations around the equilibrium cell diameter $l_0$. Computing the value of the second derivative at $l_0$ provides a value for the spring constant $k = 3(\gamma_a + \gamma_b)$. Note that we discarded potential line tensions in this analysis, although this would not qualitatively affect the results as we are considering only small deviations from mechanical equilibrium.

Given the reported role in ERK/MAPK in regulating the actomyosin cytoskeleton [8, 9], one can in principle write a dependency of any tension $\gamma_i$ on ERK activity. This means that the equilibrium length of a given cell will generically depend on ERK activity $l_0 = l_0(ERK)$. Recent experiments have indeed shown that varying ERK activity causes relative changes in F-Actin intensity between lateral and basal surfaces of MDCK cells [9]. We will next consider mechanical couplings between cells in monolayers and the interaction of ERK with mechanics. We do so for monolayers with potentially different biochemical activities and mechanical properties, to account for differences in the spatio-temporal dynamics of ERK activity present in different biological settings.

#### B. Overdamped chain of mechano-chemical springs

A major function of epithelial tissues is to form a tight barriers, which is achieved through strong mechanical couplings between neighbouring cells mediated by adhesion molecules such as E-Cadherin [10]. We write down a force balance equation for a cohesive 1D chain of epithelial cells, with cell $i$ characterized by ERK activity $E_i(t)$ and delimited by vertices $r_i(t)$ and $r_{i+1}(t)$. For vertex $r_i$, which is under frictional contact with a substrate (modelled as fluid friction with coefficient $\zeta$), this reads

$$\zeta\left(r_i(t+dt)-r_i(t)\right)=-k_i\left(r_i(t)-r_{i-1}(t)-l_i^0(t)\right)+k_{i+1}\left(r_{i+1}(t)-r_i(t)-l_{i+1}^0(t)\right) \tag{S5}$$

where $l_i^0$ and $k_i$ are the rest lengths and spring constants of a single cell derived above. In the following, we will look only up to linear stability of the model so we discard the spatial variation in $k_i$ which manifests as non-linear terms. In the continuum limit the vertex model becomes

$$\tau_r\partial_t r=\partial_{xx}r-\partial_x l^0. \tag{S6}$$

Here, lengths have been scaled by a characteristic cell length $< l >$ and we have defined a characteristic mechanical timescale $\tau_r=\zeta/k$. This timescale defines how quickly a mechanical stress diffuses in the monolayer - infinitely fast in the limiting case of vanishing friction, $\zeta\to 0$.

We now incorporate mechanosensation and response involving the chemical species ERK:

$$\begin{cases}\tau_r\partial_t r=\partial_{xx}r-\partial_x l_0\\ \tau_l\partial_t l_0=-(l_0-l_0^{eq})-\alpha(E-E_{eq})\\ \tau_E\partial_t E=-(E-E_{eq})+\beta\partial_x r\end{cases} \tag{S7}$$

The second equation represents the tendency of the rest length towards an intrinsic equilibrium cell length and the action of ERK on decreasing rest length. The third equation represents decay of ERK towards an equilibrium activity, $E_{eq}$, and the action of increasing cell length ($l=\partial_x r\leftarrow r_{i+1}-r_i$) on increasing ERK activation. Since ERK and rest lengths are carried within cells we could also write advection terms on $l_0$ and $E$ with cell velocity $\partial_t r$. However, these terms would not affect the pattern forming behaviour that we wish to study since the linear stability around the homogeneous equilibrium state is unchanged. Alternatively, dropping advection terms is equivalent to assuming that waves of ERK and strain propagate fast compared to cell velocity which also seems to hold for this system [8, 9]. Before proceeding it will help us to rewrite the model using the change of variables $l_0 \leftarrow l_0 - l_0^{eq}$ and $E \leftarrow E - E_{eq}$:

$$\begin{cases} \tau_r \partial_t r = \partial_{xx} r - \partial_x l_0 \\ \tau_l \partial_t l_0 = -l_0 - \alpha E \\ \tau_E \partial_t E = -E + \beta \partial_x r \end{cases} \tag{S8}$$

Linear stability around the homogeneous fixed state then yields the following dispersion relation:

$$(\tau_l \omega + 1)(\tau_E \omega + 1)(\tau_r \omega + q^2) = -\alpha\beta q^2 \tag{S9}$$

From this we predict a patterning instability at the critical point defined by $Re[w] = 0$ and $\frac{dRe[w]}{dq} = 0$, which translates to:

$$\alpha\beta > (\alpha\beta)_{\text{crit}} = \frac{(\tau_E + \tau_l)(\sqrt{\tau_E} + \sqrt{\tau_l})^2}{\tau_E \tau_l} \tag{S10}$$

The real and imaginary parts of $\omega$ at the critical point then define the wavenumber and angular frequency of the predicted instability, which read:

$$q_c{}^2 = \frac{\tau_r}{\sqrt{\tau_E \tau_l}}, \qquad \omega_c{}^2 = \frac{1}{\tau_l{}^{1/2} \tau_E{}^{3/2}} + \frac{1}{\tau_l{}^{3/2} \tau_E{}^{1/2}} + \frac{1}{\tau_l \tau_E}. \tag{S11}$$

Interestingly, both the wavelength and period of the instability are independent of the strength of the couplings $\alpha$ and $\beta$. Instead, they involve only the three time scales of the problem, which can all be measured from macroscopic observation. One might also notice that the period depends only on $\tau_l$ and $\tau_E$ and not on $\tau_r$. For $\alpha\beta > 0$ this is expected since the system is just a classical activator-inhibitor with delay which generically produces temporal oscillations even in the absence of spatial couplings. The mechanical time scale $\tau_r$ enters instead in the wavelength of the instability (see Fig. S2a). Its role is clearest when considering the limiting case (valid experimentally as described below) of $\tau_E \ll \tau_l$. In this limit, the period of the instability scales as $T \propto \tau_E^{1/4} \tau_l^{3/4}$ so that the wavelength scales as

$$\lambda \propto \frac{T}{\sqrt{\tau_l \tau_r}}. \tag{S12}$$

Here $\sqrt{\tau_l \tau_r}$ is the characteristic timescale of mechanical signal propagation upon biochemical change, so that the length scale of the instability is determined by how far a signal can propagate during a single temporal period of the mechano-chemical oscillation between ERK and cellular shape.

Although the linear instability threshold depends only on the product of mechano-chemical couplings $\alpha\beta$, the amplitude of mechanical vs. chemical fluctuations does depend on the relative values of couplings. We simulate the dynamics of the 1D system above the critical point ($\alpha\beta = 1.2 \times (\alpha\beta)_{\text{crit}}$) for different ratios $\alpha/\beta = 0.2, 1, 5$. As anticipated, cell shape is nearly un-affected for very small $\alpha$ while ERK levels vary drastically. In the reverse case of large $\alpha$, small ERK oscillations are accompanied by major lattice deformations and cellular stretching (Fig. S2b). Note that for numerical simulations of the model, we have to specify a non-linearity in the problem to prevent infinite activation of ERK. For this we took the simplest leading-order expansion:

$$\tau_E \partial_t E = -E - E^3 + \beta \partial_x r. \tag{S13}$$

although other non-linearities (for instance on the rest length equation $l_0$) have similar consequences.

#### C. Active migration and direction sensing

Although the above theory reproduces ERK/density waves at an appropriate wavelength and period (see Main Text), the appearance of persistent directed waves remains to be understood theoretically, together with the biophysical mechanisms through which they might guide long-range active migration [9].

To address this, we define a cell polarity $p$, which controls the direction and magnitude of active traction forces on the substrate (and thus the direction of active migration), and explore the influence of a coupling between local cell polarity $p$ and gradients of stress $\partial_x \sigma_{xx} = -\partial_x(\partial_x r - l_0)$, which is based on experimental evidence in multiple systems [13–17] (see Section I D 3 for a discussion of alternative couplings). This gives us the following model with $\zeta_0$ controlling the strength of active migration and $\gamma$ the strength of coupling between gradients of stress and ERK:

$$\begin{cases} \tau_r \partial_t r = \partial_{xx} r - \partial_x l_0 + \zeta_0 p \\ \tau_l \partial_t l_0 = -l_0 - \alpha E \\ \tau_E \partial_t E = -E + \beta \partial_x r \\ \tau_p \partial_t p = -p + D_p \partial_{xx} p + \gamma \partial_x \sigma_{xx} \end{cases} \tag{S14}$$

This model still requires a non-linearity to break symmetry of wave propagation/cell migration directionality: without it polarity oscillates around mean zero such that there is no long range order and no persistent migration. In the main text we discuss experimental evidence that suggests traction forces are a decreasing function of ERK activity and use this to motivate a non-linearity on the coupling between polarity and gradients of stress $\gamma = \gamma(E)$ [9]. This addition allows phase differences between ERK and stress (which naturally exist and are fully generic to our mechano-chemical patterns) to control the degree of symmetry breaking (see Fig. 3c for a schematic).

We analyse this in more detail below by assuming a simple saturating non-linearity of the form $\gamma(E) = \gamma/(1 + e^E)$, although all of our conclusions only require a variation of $\gamma$ with $E$. It is useful to non-dimensionalize by $T = \tau_l$ and $L = a\sqrt{\tau_l/\tau_r}$, make the change of variables $E \leftarrow \alpha E$, $l_0 \leftarrow Ll_0$ and $p \leftarrow (L/\gamma)p$, and absorb parameters $\zeta_0 \leftarrow \gamma\zeta_0/a$, $\beta \leftarrow \alpha\beta$, $D_p \leftarrow D_p/L^2$, $\tau_E \leftarrow \tau_E/T$ and $\tau_p \leftarrow \tau_p/T$ so that the model reads

$$\begin{cases} \partial_t \sigma = \partial_{xx}\sigma - \zeta_0 \partial_x p - l_0 - E \\ \partial_t l_0 = -l_0 - E \\ \tau_E \partial_t E = -E + \beta \partial_x r \\ \tau_p \partial_t p = -p + D_p \partial_{xx} p + \frac{1}{1+e^{\frac{E}{\alpha}}} \partial_x \sigma. \end{cases} \tag{S15}$$

To gain insight into this system we first derive analytical solutions for the response of an initially un-polarized tissue to an applied ERK wave (as performed for instance in Ref. [8] - although note that the theory in this study neglected polarity, set $\tau_l \to 0$ and had a different sign $\alpha < 0$ for the length-ERK coupling to the one revealed by our optogenetic experiments, see Fig. 2c-f). We impose a right-left traveling ERK wave $E(x,t) = \cos(kx + \omega t)$ and assume that the term $\zeta_0 p$ can be neglected (an approximation which renders analytical expressions more tractable, and which we verify/relax in numerical simulations of the phase diagram of Fig. 3). We also neglect diffusion of polarity, which would just act to soften oscillations without qualitatively changing the results, leaving us with a relatively simple model which can be solved analytically:

$$\begin{cases} \partial_t \sigma = \partial_{xx}\sigma - l_0 - E \\ \partial_t l_0 = -l_0 - E \\ \tau_p \partial_t p = -p + \frac{1}{1+e^{\frac{E}{\alpha}}} \partial_x \sigma. \end{cases} \tag{S16}$$

For $l_0(x,t)$ and $\sigma(x,t)$ we find sinusoidal solutions:

$$l_0\left(x,t\right) = \frac{1}{\sqrt{1+\omega^2}}\cos\left(kx+\omega t+\pi-\operatorname{atan}\left(\omega\right)\right) \tag{S17}$$

$$\sigma\left(x,t\right) = A_\sigma\cos\left(kx+\omega t+\phi_\sigma\right) \tag{S18}$$

with $A_\sigma$ and $\phi_\sigma$ given by

$$A_\sigma = \frac{1}{\sqrt{\left(1+\omega^2\right)\left(1+\frac{k^4}{\omega^2}\right)}} \qquad \phi_\sigma = \pi+\operatorname{atan}\left(\frac{k^2-\omega^2}{\omega\left(1+k^2\right)}\right) \tag{S19}$$

If we assume that $\gamma(E)$ is far from saturation we can use a linear approximation $\gamma(E) = \gamma(1/2 - E/(4\alpha))$ to obtain a solution also for the polarity:

$$\begin{aligned} p(x,t) = \frac{kA_\sigma}{8\alpha}\Big(&\sin(\phi_\sigma) + \frac{4\alpha}{\sqrt{1+\tau_p^2 w^2}}\cos\left[\omega t+kx+\phi_\sigma+\operatorname{atan}\left(\frac{1}{\tau_p\omega}\right)\right] \\ &- \frac{1}{\sqrt{1+4\tau_p^2 w^2}}\cos\left[2\omega t+2kx+\phi_\sigma+\operatorname{atan}\left(\frac{1}{2\tau_p\omega}\right)\right]\Big) \end{aligned} \tag{S20}$$

From Eq. S20, we see that symmetry is broken by the appearance of a constant term and the shape of the oscillation is altered by the appearance of a phase-shifted second harmonic. The constant term is also the average polarity $\overline{p}$ which is approached in the limit $\tau_p \to \infty$:

$$\overline{p}(\omega,k) = \lim_{\tau_p\to\infty} p(x,t|\omega,k) = \frac{kA_\sigma(\omega,k)}{8\alpha}\sin\left(\phi_\sigma(\omega,k)\right) = \frac{1}{8\alpha}\frac{\omega k\left(\omega^2-k^2\right)}{\left(1+\omega^2\right)\left(k^4+\omega^2\right)} \tag{S21}$$

Thus, the average polarity is determined by the angular frequency $\omega$ and wavenumber $k$ of the applied wave (plotted in Fig. S5a). From Eq. S21 we also see that there is a boundary between net-positive and net-negative polarization at $\omega = k$, and that polarity is maximized in the direction opposite to the ERK/density waves for a unique value of the wavelength and period, which corresponds to a phase difference $\phi_\sigma = 5\pi/4$ between stress and ERK (with the coefficient $\alpha$ changing only the value of the optimum and not its position). Analysis gives expressions for the optimal wavelength and period:

$$k = 1 \text{ and } \omega = 1+\sqrt{2} \tag{S22}$$

or adding back dimensions,

$$\lambda = 2\pi\sqrt{\frac{\tau_l}{\tau_r}} \text{ (in units of cell length)} \quad \text{and} \quad T = \frac{2\pi}{1+\sqrt{2}}\tau_l \tag{S23}$$

With experimentally estimated values of $\tau_r$ and $\tau_l$ (see Fig. 2 and note that $\tau_E$ does not appear because we are enforcing the dynamics on ERK) we predict optimal waves at wavelengths of $15-55$

cells and periods of $2-10$ hours (Fig. S5a). However, this analytical criteria does not take into account ERK de-sensitisation, occurring on time scales of roughly an hour, which manifests in slow, long-term, decays of ERK activity following both mechanical stretch (Fig. 2b) and optogenetic ERK activation experiments [9]. We anticipate that this effect will decrease sensitivity to long-period oscillations and when we include it in numerical simulation of the full model ($\tau_d$ = 1h, see following section Section I D 1 for further theoretical details), we see that the optimum moves towards shorter periods and wavelengths (albeit relatively little, displaying robustness of the results, see Fig. 3d and Fig. S5). The predicted optimum then occurs at wavelengths of $\lambda \approx 15-40$ cells and periods of $T \approx 2-8$ hours.

Strikingly, this range is close to the values observed in multiple systems, including the MDCK cells which we study:

- In our conditions, and Ref. [8, 9]: $\lambda \approx 10-20$ cells, $T \approx 1-2$ h,
- In Ref. [12]: $\lambda \approx 50$ cells (our estimation from their report of a wavelength around 1mm and cell diameter of 20$\mu$m), $T \approx 2$ h
- in Ref. [19]: $\lambda \approx 18$ cells (our estimation from their report of a wavelength of $370\mu m$ and cell diameter of 20$\mu$m), $T \approx 4.7$ h
- in Ref. [20]: $T \approx 6$h, and a wavelength of at least 700$\mu$m (as cells oscillate coherently on a 700$\mu$m diameter disk), i.e. approx 35 cells
- From figures in Ref. [21]: we estimate a period $T \approx 5$h and a wavelength of $\lambda \approx 350\mu$m (approx. 15-20 cells)
- From Ref. [22]: $T \approx 2.6$h and a wavelength of $\lambda \approx 15$ cells (estimated combining their measurement of period with their estimation of propagation speed of the wave, leading to around 300$\mu$m).
- In Ref. [23], which examines *in vivo* mouse skin upon wound healing, we went back to the data and estimated $\lambda \approx 8$ cells, $T \approx 1.3$ h

We note that some discrepancies in the wavelength and period observed for MDCK cells are likely to be due to differences in experimental set-ups. Different substrates for instance are expected to modify the friction coefficient $\zeta$ of monolayers ($\tau_r = \zeta/k$), thus directly impacting the wavelength of the instability, as well as potentially affecting the time scales of ERK signalling ($\tau_E$ and $\tau_l$) via mechano-sensation. However, it remains striking that our measurements, as well as published values of wavelength and period, indicate that the system is close to optimality in terms of polarization strength in response to a mechano-chemical wave.

Furthermore, although we have so far considered the limiting case of driven ERK dynamics (in which any wave can be applied on the system), the biophysical origin of the instability places additional constraints on the system. In particular, it restricts the period and wavelength of the instability to a curve in $\omega - k$ space (i.e. the dispersion relation of our instability, see Fig. 3d or Fig. S5). Interestingly, we find that the emergent waves possible from our biophysical model have dominant modes governed by $w^4 = k^4 + k^3 + k^2$ such that they can only induce polarization in the direction opposite to ERK propagation (i.e. sitting in the side of the diagram where $w > k$), for any values of model parameters! This makes it an extremely robust mechanism for coordinating cell migration in a unidirectional manner.

#### D. Sensitivity to model assumptions

In these section, we discuss different extensions to our model, as well as alternative couplings, to provide insights as to its robustness.

##### *1. Adaptation of ERK to mechanical stresses*

Our model for ERK dynamics (Eq. S8) can be extended to account for slow adaptation of ERK in response to constant stretch or long-term optogenetic activation ([9] and Fig. 2). In its simplest form, a model of long-term ERK adapation reads:

$$\tau_E \partial_t^2 E + \partial_t E = -E/\tau_d + \beta \partial_t \partial_x r. \qquad \text{(S24)}$$

as can be seen from the limiting case of this equation: on time-scales longer than $\tau_d$, ERK relaxes towards zero (irrespective of cell area), whereas on time-scales shorter than $\tau_d$, we recover the previous behavior, with ERK linearly following the deformation $\partial_x r$ with a delay $\tau_E$. One can then recalculate the dispersion relation for this extended/adaptive model:

$$(\tau_E w^2 + w + 1/\tau_d)(\tau_l w + 1)(\tau_r w + q^2) = -\alpha\beta w q^2 \qquad \text{(S25)}$$

As expected this reduces to the previous dispersion relation for $\tau_d \to \infty$. We estimated a lower bound on $\tau_d$ by fitting mechanical stretch experiments (Fig. 2a,b) with numerical solutions of Eq.

S24 for an instantaneously applied constant stretch (temporal step function in $\partial_x r$ - see Fig. S3c) and estimate $\tau_d \approx 1$h. Importantly, the dispersion relation for corresponding values of $\tau_d$ shows only slight differences to the dispersion relation (even for short de-sensitization time scales of $\tau_d = 30$min, Fig. S3d), arguing that adaptation has a negligible effect on pattern formation in this system. One should note however that adaptation should be important for longer-term expansion experiments on time scale of days, although in this case other effects such as cell division [11, 12] also become important. We also discussed above (Section I C) how adaptation slightly reduces the effectiveness of very-long period (and wavelength) waves for inducing strong polarization (see Fig. S5).

#### 2. *Direct cell-cell biochemical communication*

In our model, ERK is restricted to the subcellular level, and does not diffuse directly from one cell to the next, instead propagating via mechanical couplings and the mechano-sensitivity of ERK (which is validated by the analysis of the experiments of Fig. 2 and [9]). An extension could consider that biochemical relays act as an effective diffusion coefficient $D_E$ appearing directly in the equation on ERK dynamics:

$$\tau_E \partial_t E = \tau_E D_E \partial_{xx} E - E + \beta \partial_x r. \tag{S26}$$

This allows for diffusive biochemical communication over the lengthscale $\sqrt{D_E \tau_E}$ and translates into a modified dispersion relation:

$$(\tau_E w + 1 + D_E q^2)(\tau_l w + 1)(\tau_r w + q^2) = -\alpha\beta w q^2. \tag{S27}$$

Plotting this dispersion relation for $D_E = 1$, a typical diffusion rate for biological proteins ($D_E \approx 10^{-11} m^2 s$ in real units for cell length $< l > \approx 20\mu m$), shows that this would have only very weak effects on the instability and pattern formation (Fig. S3b).

#### 3. *Alternative coupling to cell polarity*

In the main text, we explore the influence of a coupling between local cell polarity $p$ and gradients of stress, which is based on experimental evidence in multiple systems [13–17]. There are, however, other proposals for how tissue mechanics impacts cellular polarity, and in particular the possibility that cellular polarity $\vec{p}$ aligns with and cellular velocity $\vec{v}$ (which are not necessarily identical in confluent monolayers due to mechanical couplings between cells [17, 18]). This would modify the equation for polarity in Eq. S14 as follows:

$$\tau_p \partial_t p = -p + D_p \partial_{xx} p + \gamma v \tag{S28}$$

Notice however that gradients of stress and cell velocity $v = \partial_t r$ are closely related by force balance:

$$\tau_r \partial_t r = -\partial_x \sigma_{xx} + \zeta_0 p \tag{S29}$$

such that re-injecting velocity into Eq. S28 returns a term on gradients of stress:

$$\tau_p \partial_t p = -(1 - \frac{\gamma \zeta_0}{\tau_r})p + D_p \partial_{xx} p - \frac{\gamma}{\tau_r} \partial_x \sigma_{xx}. \tag{S30}$$

This term is similar, up to a rescaling factor, to the coupling assumed in main text but with opposite sign. This sign difference is important because it reverses the direction in which cells polarize in response to ERK waves, i.e. polarity-velocity coupling produces migration in the opposite direction compared to the model in the main text (stable only as long as $\gamma \zeta_0 < 1$.) All of the conclusions regarding optimality are therefore reversed and emergent waves robustly produce polarity in the "wrong" direction, leading us to favour our original coupling. Finally, considering a direct polarity alignment between cells (modelled as a diffusion coefficient $D_p$ in the polarity equation), does not change qualitatively any of the results discussed in the main text so we omit it in simulations.

### II. NUMERICAL SIMULATIONS

To simulate our system in 1D, we integrate the discrete equations of the chain-of-springs model using a simple Euler method. We use a time step of numerical integration $\delta t = 0.25$min which is an order of magnitude smaller than any other time scale in the problem. The length scale is set by the average homeostatic cell length $< l >= 1$, with spatial derivatives taken as difference to the nearest neighbour. We simulate a chain of N cells, with labels $i$, ERK activity $E_i$, preferred length $l_0^i$, length $l_i = r_{i+1} - r_i$ and polarity $p_i$.

To simulate confluent tissues, we used periodic boundary conditions. To simulate monolayer expansion, we first simulate the system for 100 min without leader cells or boundary expansion before release (to allow for ERK dynamics to equilibrate), and complemented the problem with a boundary condition at the free edge such that cell polarity approached $p_N = p_b$. This provides a fixed polarity to leader cells at the edge of the monolayer (as observed experimentally, see Ref.

[24]). We also applied small amounts of white noise of amplitude $\eta_E = 0.05$ to the equation of ERK dynamics.

To simulate optogenetically applied ERK waves of frequency $\omega$ and wavevector $q$ and construct Fig. 3d, we imposed traveling waves $E(x,t) = E_0 \cos(\omega t - qx)$ and monitored the average response of other variables (in particular cellular displacement) after 10 temporal periods of oscillation. In order to find a simple analytical solution for average polarity in the presence of an applied ERK wave, and construct Fig. 3e and Fig. S5a, we made several assumptions: we neglect traction forces and diffusion of polarity, as well as advection of ERK, rest length and polarity. In contrast, we include all of these effects in numerical simulations of applied waves and of the fully coupled system (Fig. 3d, Fig. S5d, Fig. 4 and Fig. S6), with the comparable analytical and numerical results (Fig. S5) showing little qualitative difference.

To simulate the experiments of uniform activation (resp. inhibition of ERK) shown in Fig. S2c, we used the same simulation as above for confluent tissues, with the only change being to add a constant term to the equation of ERK, which then reads

$$\tau_E \partial_t E = E_0 - E - E^3 + \beta \partial_x r. \tag{S31}$$

Given that ERK levels in the control simulation oscillates between the range of $-0.15$ and $+0.15$, we set $E_0 = 0.25$ (resp. $E_0 = -0.25$) to simulate global ERK activation (resp. inhibition) and plot the corresponding dynamics for local ERK activity and cell length (Fig. S2c). This demonstrates a marked inhibition of density waves in both conditions, in agreement with experimental data (Fig. S1).

### III. EXPERIMENTS

#### A. Cell culture

MDCK cells (RIKEN BRC, RCB0995) were used for this study. All cell experiments were performed with a culture medium including Medium 199 (11043023; Life Technologies, Carlsbad, CA), 10% FBS (no. 172012-500ML, SIGMA, St. Louis, MO), 100unit mL$^{-1}$ penicillin, and 100$\mu$g.mL$^{-1}$ streptomycin (no. 26253-84, Nacalai, Kyoto, Japan). The details were described elsewhere [9].

#### B. Cell migration assay

For the confinement assay, the cells expressing EKAREV-NLS [25] were seeded inside a culture insert (ibidi, Martinsried, Germany) ($7 \times 10^3$ cell/well), and cultured for 24 hours. The cells were then released and imaged with an epifluorescence microscope at intervals of 5 min beginning 30 min after an exchange of culture media [9]. For the migration assay with high density, cells were cultured for a longer period of 48 hours prior to release to increase the cell number.

#### C. CRISPR/Cas9-mediated knockout of Merlin

LentiCRISPRv2-bleo was constructed by replacing puromycin resistance gene in lentiCRISPRv2 (Addgene Plasmid: no. 52961) with bleomycin resistance one. For CRISPR/Cas9-mediated KO of dog NF2 (Merlin), two single guide RNAs (sgRNA) targeting the exons of NF2 were designed using the CRISPRdirect [26]. The following sequences were used for the sgRNA sequences: CCTGGCTTCTTACGCCGTCC (sgRNA1) and GACCCCTCTGTTCACAAACG (sgRNA2). Oligo DNAs for the sgRNA1 and sgRNA2 were cloned into the lentiCRISPRv2 and lentiCRISPRv2-bleo vectors, respectively. To increase the KO efficiency, the sgRNA1 and sgRNA2 together with Cas9 were simultaneously introduced into MDCK cells by lentivirus. The infected cells were selected with $2.0\mu$g.mL$^{-1}$ puromycin and $100\mu$ g.mL$^{-1}$ zeocin. After the selection, reduction in expression levels of the proteins was confirmed by immunoblotting. Bulk cells were used for the experiments.

Statistical analyses between Merlin and WT patterns were performed with MATLAB. No statistical analysis was used to predetermine the sample size. The sample sizes, statistical tests, and p-values are indicated in the figure legends. To compare two sets of data, paired t-tests were used. P-values of less than 0.05 were considered to be statistically significant in two-tailed tests.

#### D. Microscopy

Details of time-lapse FRET imaging and traction force microscopy were described elsewhere [9].

### IV. FITTING OF MODEL PARAMETERS AGAINST EXPERIMENTAL DATA

A major assumption of our model is the bidirectional coupling between ERK activity and cellular shape, characterized by time scales $\tau_E$ and $\tau_l$ and coupling strengths $\alpha$ and $\beta$. Ideal experiments for constraining these parameters were performed in Ref. [9]: they consist of i) applying abrupt mechanical strains while observing ERK response (mechanical stretching experiments combined with FRET sensors, Fig. 2a), and conversely ii) applying abrupt changes in ERK activity while observing mechanical response (optogenetic ERK activation experiments, Fig. 2c). Below, we detail our theoretical interpretation and statistical analysis of each of these experiments.

#### A. Mechanical stretching experiments

The cellular stretching experiment is relatively straightforward to interpret from a theoretical perspective since it involves only the equation on ERK (See Eq. S8). At time $t = 0$ an abrupt, uniform and permanent strain $\Delta\epsilon = 50\%$ is applied to the cellular monolayer. We model this as a Heaviside stretch $\Delta\epsilon H(t)$:

$$\tau_E \partial_t E = -E + \beta \Delta\epsilon H(t) \tag{S32}$$

which predicts that ERK follows

$$E(t) = \beta \Delta\epsilon (1 - e^{-t/\tau_E}), \tag{S33}$$

and allows us to use the timescale of ERK activation to infer $\tau_E$ and the magnitude of the response to infer $\beta$ (see Fig. 2b and Fig. S4a). We examined average ERK activity in three independent repetitions of the stretch experiment (at least n = 685 averaged cells in each), and found that good fits to the data could be achieved for $\tau_E = 6 \pm 2$ min, $\beta = 0.56 \pm 0.06$ and $E_{eq} = 0.66 \pm 0.05$ (mean $\pm$ standard deviation). Fitting was performed using a standard gradient descent routine (BFGS) from SciPy optimize with least square error function. We note that the applied stretch is larger than the typical stretch experienced by cells during normal oscillations, which means that we likely underestimate $\beta$, due to potential saturation effects at large areas.

#### B. Optogenetic experiments

The optogenetic ERK activation experiment is slightly more complicated to investigate as it involves both the equations on cell displacement and rest length (see Eq. S8). When ERK is activated in a half-plane, the system is symmetric parallel to the boundary, so we model in onedimension with a generic Heaviside function for the optogenetic ERK activation:

$$\tau_r \partial_t r = \partial_{xx} r - \partial_x l_0 \tag{S34a}$$

$$\tau_l \partial_t l_0 = -l_0 - \alpha(\Delta E H(x) + E_0) \tag{S34b}$$

which after non-dimensionalization using $T = \tau_l$ and $L = a\sqrt{\tau_l/\tau_r}$ reads

$$\partial_t r = \partial_{xx} r - \sqrt{\frac{\tau_l}{\tau_r}} \partial_x l_0 \tag{S35a}$$

$$\partial_t l_0 = -l_0 - \alpha\sqrt{\frac{\tau_r}{\tau_l}}(\Delta E H(x) + E_0). \tag{S35b}$$

The second equation is solved independently to give $l_0(x,t) = \alpha\sqrt{\frac{\tau_r}{\tau_l}}(e^{-t} - 1)(\Delta E H(x) + E_0)$ which we substitute into the equation on $r$ to give

$$\partial_t r = \partial_{xx} r + \alpha \Delta E(1 - e^{-t})\delta(x) \tag{S36}$$

This describes diffusion from an exponential point source, but can be reformulated as diffusion from a constant point source on new variable $m(x,t) = r(x,t) + \partial_t r(x,t)$:

$$\partial_t m = \partial_t(r + \partial_t r) = \partial_{xx} m + \alpha \Delta E \delta(x) \tag{S37}$$

We can solve this equation for $m(x,t)$ either in Fourier space or by integrating the solution for an instantaneous point source as follows:

$$m(x,t) = \int_0^t \frac{\alpha \Delta E}{4\pi\tau} \exp\left[\frac{-x^2}{4\tau}\right] d\tau = \alpha \Delta E \left( \sqrt{\frac{t}{\pi}} \exp\left[\frac{-x^2}{4t}\right] + \frac{x}{2} \operatorname{erf}\left[\frac{x}{2\sqrt{t}}\right] - \frac{|x|}{2} \right) \tag{S38}$$

Finally, substituting this solution back into the definition of $m(x,t)$ solving for $r(x,t)$ gives a result for the displacement,

$$r(x,t) = \frac{\alpha \Delta E}{2} \left( |x| \left( \operatorname{erf}\left[\frac{|x|}{2\sqrt{t}}\right] - 1 \right) + e^{-t} \left( -ie^{-i|x|} + 2\sqrt{\frac{t}{\pi}} \exp\left[t - \frac{x^2}{4t}\right] - e^{ix} \operatorname{erfi}\left[\sqrt{t} - \frac{ix}{2\sqrt{t}}\right] \right) \right) \tag{S39}$$

and velocity,

$$\partial_t r(x,t) = \frac{\alpha \Delta E}{2} e^{-t} \left( ie^{i|x|} + e^{ix} \operatorname{erfi}\left[\sqrt{t} - \frac{ix}{2\sqrt{t}}\right] \right) \tag{S40}$$

Note that $r(x,t)$ describes the field of cellular deformations - properly interpreted as the displacement after time $t$ of a cell that starts a distance $x$ from the boundary - and we tracked this throughout the experiment in order to fit the solutions. The solution for the boundary ($x = 0$) is much simpler than for the bulk:

$$m(0,t) = \frac{\alpha \Delta E}{\sqrt{\pi}} t^{1/2} \tag{S41}$$

$$r(0,t) = \frac{\alpha \Delta E}{\sqrt{\pi}} t^{1/2} - \frac{\alpha \Delta E}{2} e^{-t} \operatorname{erfi}\left(t^{1/2}\right) \tag{S42}$$

$$\partial_t r(0,t) \equiv v(0,t) = \frac{\alpha \Delta E}{2} e^{-t} \operatorname{erfi}\left(t^{1/2}\right) \tag{S43}$$

and because the initial displacement is controlled by contraction of only the first cell layer we were able to use the power law on $m(0,t)$ to estimate $\tau_l = 100 - 140$min (Fig. 2d,e). When dimensions are added back the power law becomes

$$Lm = r + \tau_l v = L \frac{\alpha \Delta E}{\sqrt{\pi}} \left(\frac{t}{\tau_l}\right)^{1/2} = \frac{a \alpha \Delta E}{\sqrt{\pi}} \left(\frac{t}{\tau_r}\right)^{1/2}, \tag{S44}$$

or, in log-log space,

$$\log(Lm) = \log(r + \tau_l v) = \log\left(\frac{a \alpha \Delta E}{\sqrt{\pi \tau_r}}\right) + \frac{1}{2}\log(t). \tag{S45}$$

Since $\tau_r$ controls how quickly deformation spreads into the bulk we instead estimate it using the full solution (Eq. S39) which gives the range $\tau_r = 4 - 11$ (Fig. 2f and Fig. S4b-d). We can also resolve $\alpha$ from estimates of $\alpha \Delta E$ (see Fig. S4b-d) using the observation that the step in optogenetic ERK activity is typically around one tenth of the initial value ($\Delta E / E_0 \approx 0.1$) to give $\alpha = 7 \pm 2$ (mean $\pm$ standard deviation). We note however that, like for our estimate of $\beta$, we likely under-estimate this value, because we neglect the active ERK response of the non-activated patch of cells that is being stretched by the activated ERK patch, which would tend to decrease the overall amplitude of contraction. This would explain why the coefficients of coupling that we infer are smaller than the predicted threshold of the instability (see Section I B).

#### C. Data analysis and fitting

The optogenetic experiments (Fig. 2c) used raw data from [9] and consisted of N=3 independent repeats, with each repeat consisting of n=3 fields of view along the expanding monolayer. To quantify the boundary displacement we averaged the displacement from initial position $r(0,t)$ across all three fields of view for each repeat and calculated velocity $v(0,t)$ using a standard NumPy routine for central differences. For the bulk we tracked the displacement of cell centers $r(x,t)$ using Imaris software and binned cells from each repeat according to initial distance from the boundary $x$. To account for the heterogeneity in cell responses, we fitted each repeat independently to report the broad range of parameter values (Fig. 2d,e and Fig. S4b-d), and also averaged all three to perform the collective bulk fitting in Fig. 2f. Where necessary we used a global minimization algorithm (basinhopping from SciPy optimize), otherwise a standard gradient descent (BFGS from the same library), with least-squares error function in all cases. We set the zero point ($t = 0$) to 10 min (i.e. two timepoints) after optogenetic activation to account for the transient in ERK activation in response to the optogenetic switch, which occurs on timescales of $\tau_E$ (i.e. to account for the lack of an instantaneous activation as assumed in the model) and discarded the next two timepoints to avoid any remaining bias from this transient. We first fit the boundary (Eq. S45) to estimate $\tau_l$ before fixing these values while estimating $\tau_r$ from the bulk (Eq. S39). For the collective fit in Fig. 2f we fixed $\tau_l$ using the average value between repeats.
